# Evolving super stimuli for real neurons using deep generative networks

**DOI:** 10.1101/516484

**Authors:** Carlos R. Ponce, Will Xiao, Peter F. Schade, Till S. Hartmann, Gabriel Kreiman, Margaret S. Livingstone

## Abstract

Finding the best stimulus for a neuron is challenging because it is impossible to test all possible stimuli. Here we used a vast, unbiased, and diverse hypothesis space encoded by a generative deep neural network model to investigate neuronal selectivity in inferotemporal cortex without making any assumptions about natural features or categories. A genetic algorithm, guided by neuronal responses, searched this space for optimal stimuli. Evolved synthetic images evoked higher firing rates than even the best natural images and revealed diagnostic features, independently of category or feature selection. This approach provides a way to investigate neural selectivity in any modality that can be represented by a neural network and challenges our understanding of neural coding in visual cortex.

**Highlights:**

- A generative deep neural network interacted with a genetic algorithm to evolve stimuli that maximized the firing of neurons in alert macaque inferotemporal and primary visual cortex.
- The evolved images activated neurons more strongly than did thousands of natural images.
- Distance in image space from the evolved images predicted responses of neurons to novel images.

## 1. Introduction

A transformative revelation in neuroscience was the realization that visual neurons respond preferentially to unique stimuli (Hubel & Wiesel, 1962). Those findings opened the doors to investigate neural coding for myriad stimulus attributes. A central challenge in elucidating neuronal tuning in visual cortex is the impossibility of testing all stimuli. Even for a small patch of 100 ∗ 100 binary pixels, there are 2^10,000^ possible images, a problem that becomes even more intractable in color. Symmetry assumptions and natural image statistics reduce the problem but it is still not feasible to present a neuron with all possible natural stimuli. Investigators circumvent this formidable empirical challenge by employing ad hoc hand-picked stimuli, inspired by hypotheses that particular cortical areas encode specific visual features (Felleman & Van Essen, 1987; Zeki, 1973, 1974). This approach has led to important insights through the discovery of cortical neurons that respond to stimuli depicting different motion directions (Hubel, 1959), color (Michael, 1978), binocular disparity (Barlow et al., 1967), curvature (Pasupathy & Connor, 1999), and even complex natural shapes such as hands or faces (Desimone et al., 1984; Gross et al., 1972).

Despite the successes using hand-picked stimuli, the field may have missed image types that better reflect the “true” tuning of cortical neurons. A series of interesting alternative approaches have addressed this question. One is to start with hand-picked stimuli that elicit strong activation and systematically deform those stimuli; this approach has revealed that neurons often tend to respond even better to distorted versions of the original stimuli (Freiwald et al., 2009; Kobatake & Tanaka, 1994; Leopold et al., 2006). Another approach involves presenting noise stimuli and averaging the stimuli that occurred just before a spike (Gaska et al., 1994; Jones & Palmer, 1987), but this has not yielded useful results in higher cortical areas, because it cannot capture non-linearities. An elegant alternative is to use a genetic algorithm whereby the neuron under study can itself dictate which stimuli it prefers. A successful implementation of this idea by Connor and colleagues (Yamane et al., 2008) investigated selectivity in macaque V4 and IT; however, this approach relied on a limited parametrized stimulus space. To investigate the tuning properties of inferior temporal cortex (IT) neurons in macaque monkeys, here we extend prior approaches by using a pre-trained deep generative neural network (Dosovitskiy & Brox, 2016) and a genetic algorithm to allow neuronal responses to direct the evolution of synthetic images. This generative network had been previously used to synthesize images that would maximally activate units in various convolutional neural networks (Nguyen et al., 2016), which emulate aspects of computation along the primate visual stream (Yamins et al., 2014). This network can synthesize high-quality, diverse images, which encompass a vast, unbiased, and explorable image space (Nguyen et al., 2016). The network takes 4096-dimensional real-valued vectors (image codes) as input and deterministically transforms them into 256×256 RGB synthetic images (METHODS, Figure 1A,D). Here a genetic algorithm used responses of neurons recorded in alert macaques to optimize image codes to this network. At the start of each experiment, the network created an initial population of 40 images from random achromatic Portilla and Simoncelli textures (Portilla & Simoncelli, 2000) (Figure 1B). We recorded responses of IT neurons (spike counts 70–200 ms after stimulus onset minus background) while monkeys engaged in a passive fixation task. Images subtended 3°×3° and covered the unit’s receptive field. Neuronal responses to each synthetic image were used to score the image codes. In each generation, new images were synthesized from the top 10 image codes, unchanged, from the previous generation plus 30 new image codes generated by firing-rate-based selection of all the codes from the preceding generation, recombination, and mutation (Figure 1D). This process was repeated for up to 250 generations over 1–3 hours; session duration depended on the monkey’s willingness to maintain fixation. To monitor potential changes in firing rate due to adaptation and to compare synthetic-image responses to natural-image responses, we interleaved reference images that included faces, body parts, places and simple line drawings. We conducted evolution experiments on IT neurons in five monkeys: two with chronic microelectrode arrays in posterior IT (PIT, monkeys Ri and Gu), two with chronic arrays in central IT (CIT, monkeys Ge and Y1), and one with a recording chamber over CIT (monkey B3). Lastly we validated the approach in a sixth monkey with a chronic array in primary visual cortex (V1, monkey Vi).

**Figure 1.**
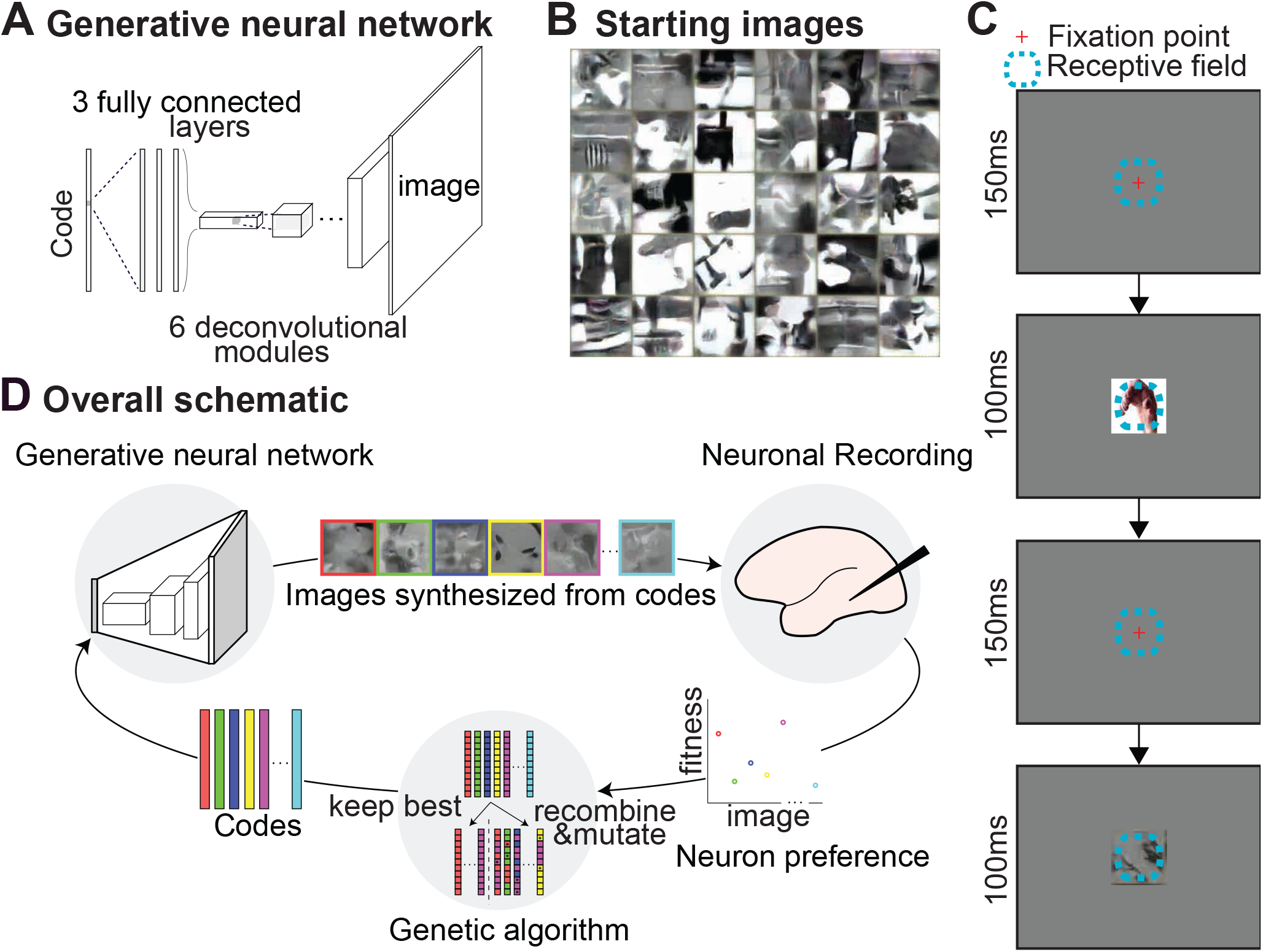
Synthesis of optimum images via neuron-guided evolution. (**A**) Generative network. Architecture of the pre-trained deep generative network (Dosovitskiy & Brox, 2016). The network comprised three fully connected layers and six deconvolutional modules. (**B**) The initial synthetic images were random achromatic Simoncelli and Portilla textures. (**C**) Behavioral task. Animals fixated within a 2.0°-diameter window while images were presented for 100 ms followed by a 100 to 200 ms blank period. Red cross: fixation point; dashed line, population RF. (**D**) Experimental flow. Image codes were forwarded through the deep generative network to synthesize images presented to the monkey.

## 2. Results

### 2.1. Evolution of synthetic images by units in CaffeNet

We first tested this approach on units in an artificial neural network, as models of biological neurons. Our method generated superstimuli for units across layers in CaffeNet, a variant of AlexNet Figure 2. The evolved images were frequently better stimuli than all of >1.4 million images that included the training set of the network. For units in the first and last layers, the method produced stimuli that matched the ground truth best stimuli in the first layer and category label in the last layer. [This method works even if we artificially inject noise into these units to better model stochastic neurons (data not shown).]

**Figure 2.**
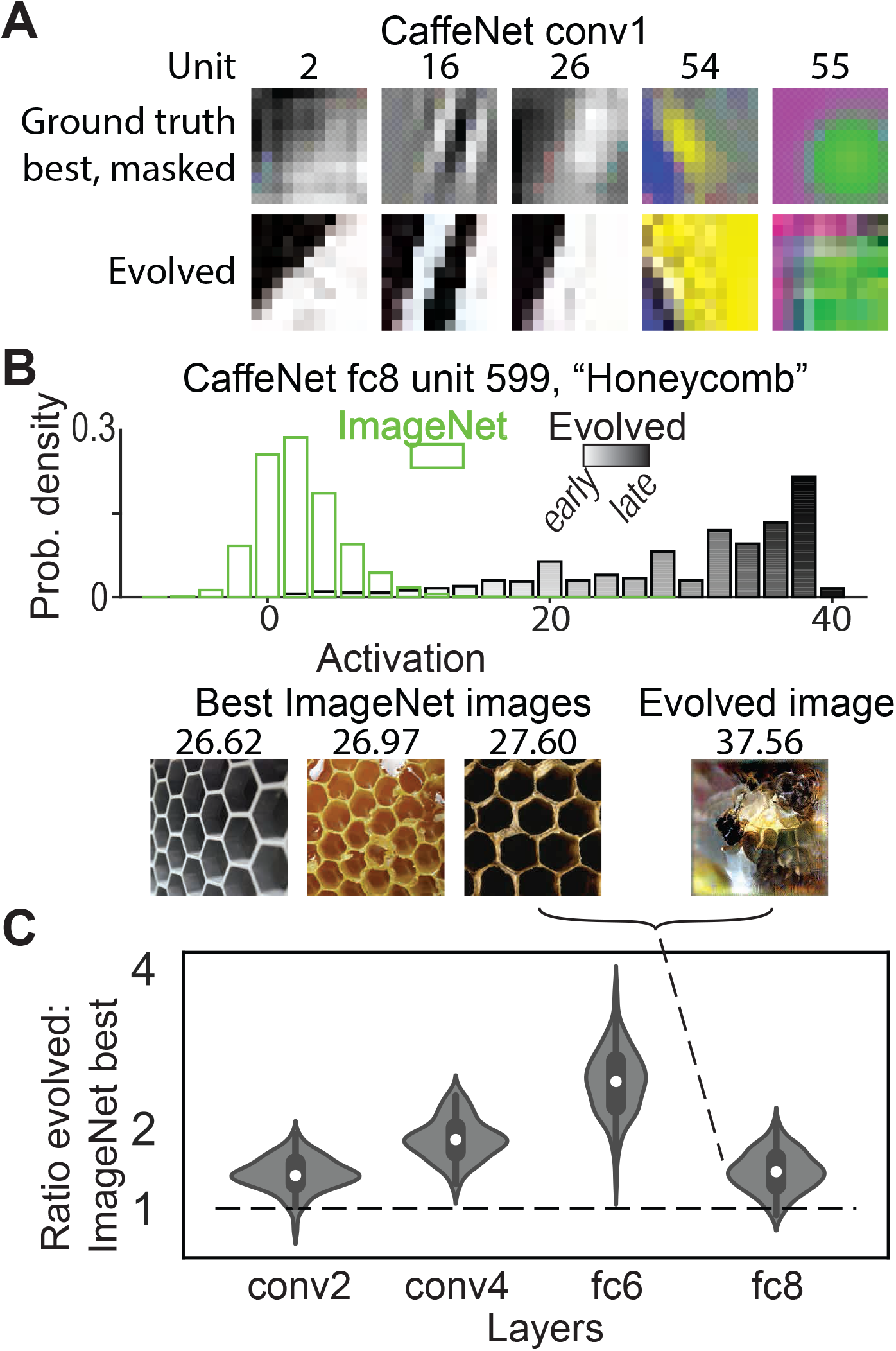
The generative evolutionary algorithm produced superstimuli for units in CaffeNet. (**A**) Evolved images resembled the ground truth best in the first layer of CaffeNet. In the ground truth best, transparency indicates the relative contribution of each pixel to the overall firing rate. Only the center 11×11 pixels of the evolved images are shown, matching the filter size of the units. (**B, C**) Most evolved images activated artificial units more strongly than all of > 1.4 million images in the ILSVRC2012 dataset (“ImageNet”, (Deng et al., 2009)). (**B**) Top, distribution of activations to ImageNet images and evolved images for one unit. Grayscale gradient indicates generation of evolved images. Bottom, best 3 ImageNet images and one evolved image labeled with their respective activations. In this case, the evolved image activated the unit ~1.4× as strongly as did the best ImageNet image. (**C**) Distribution of (evolved:best in ImageNet) ratios across 4 layers in CaffeNet, 100 random units each layer.

### 2.2. Evolution of synthetic images by a single biological neuron

We first show an example of an evolution experiment for one PIT single unit (Ri-10) in chronic-array monkey Ri. The synthetic images changed with each generation as the genetic algorithm optimized the images according to the neuron’s responses (Figure 3). At the beginning of the experiment, this unit responded more strongly to the reference images than to the synthetic images, but over several generations, the synthetic images evolved to become more effective stimuli (Figure 4A). To quantify the change in responses over time, we fit an exponential function to the cell’s firing rate as a function of generation number (solid thick lines in Figure 4A). This neuron showed an increase in response rate of 51.5±5.0 (95% CI) spikes per s in response to the synthetic images and a decrease of −15.5±3.5 spikes per s to the reference images (see Table S1 for quantification across monkeys) — thus the synthetic images became gradually more effective, despite the neuron’s slight reduction in firing rate to the reference images, presumably due to adaptation.

**Figure 3.**
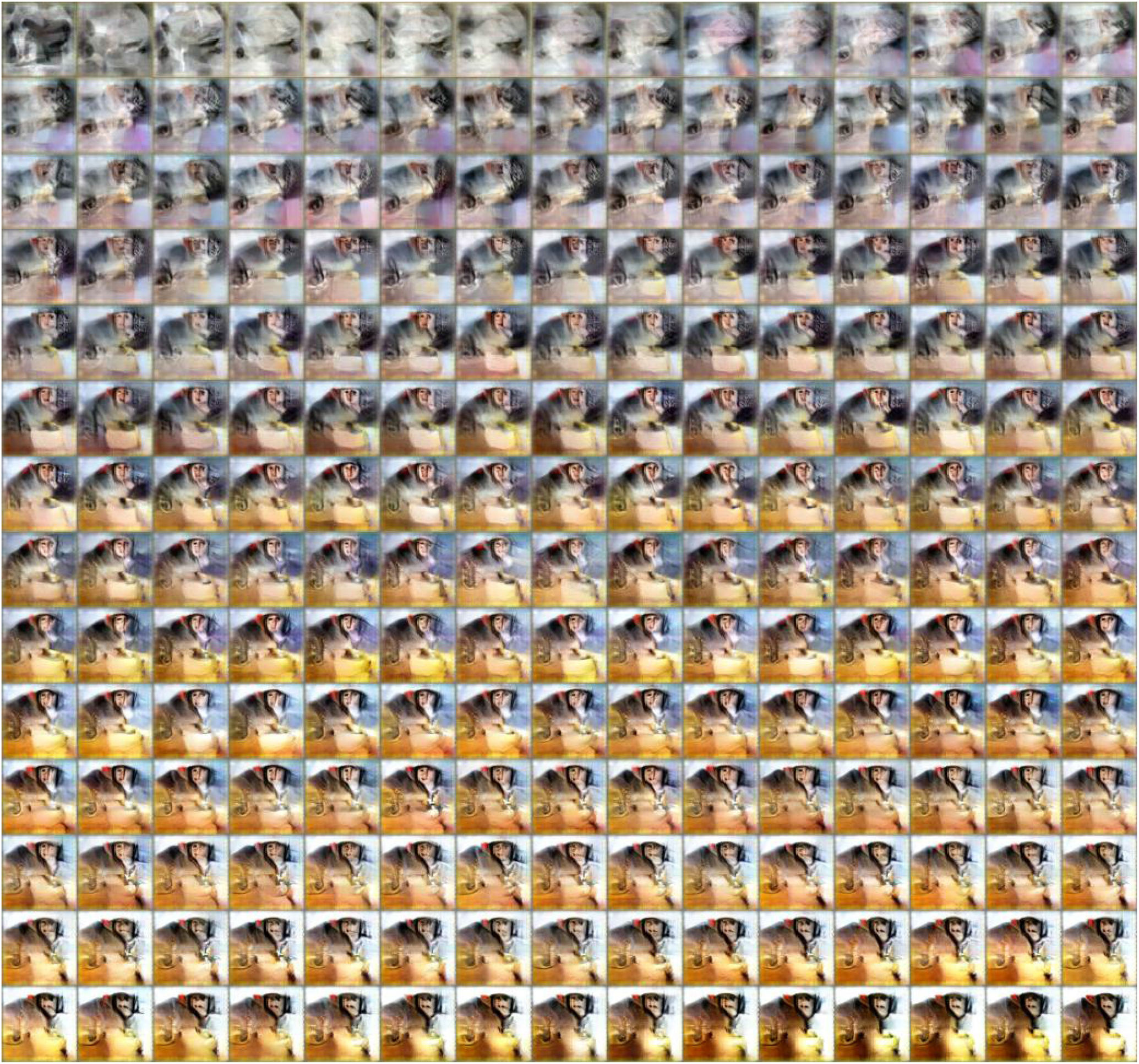
Evolution of synthetic images by a single monkey-selective neuron, Ri-10. Each image is the average of the top 5 synthetic images for each generation (ordered L to R, top to bottom). The response of this neuron in each of these generations is shown in Figure 4A.

**Figure 4.**
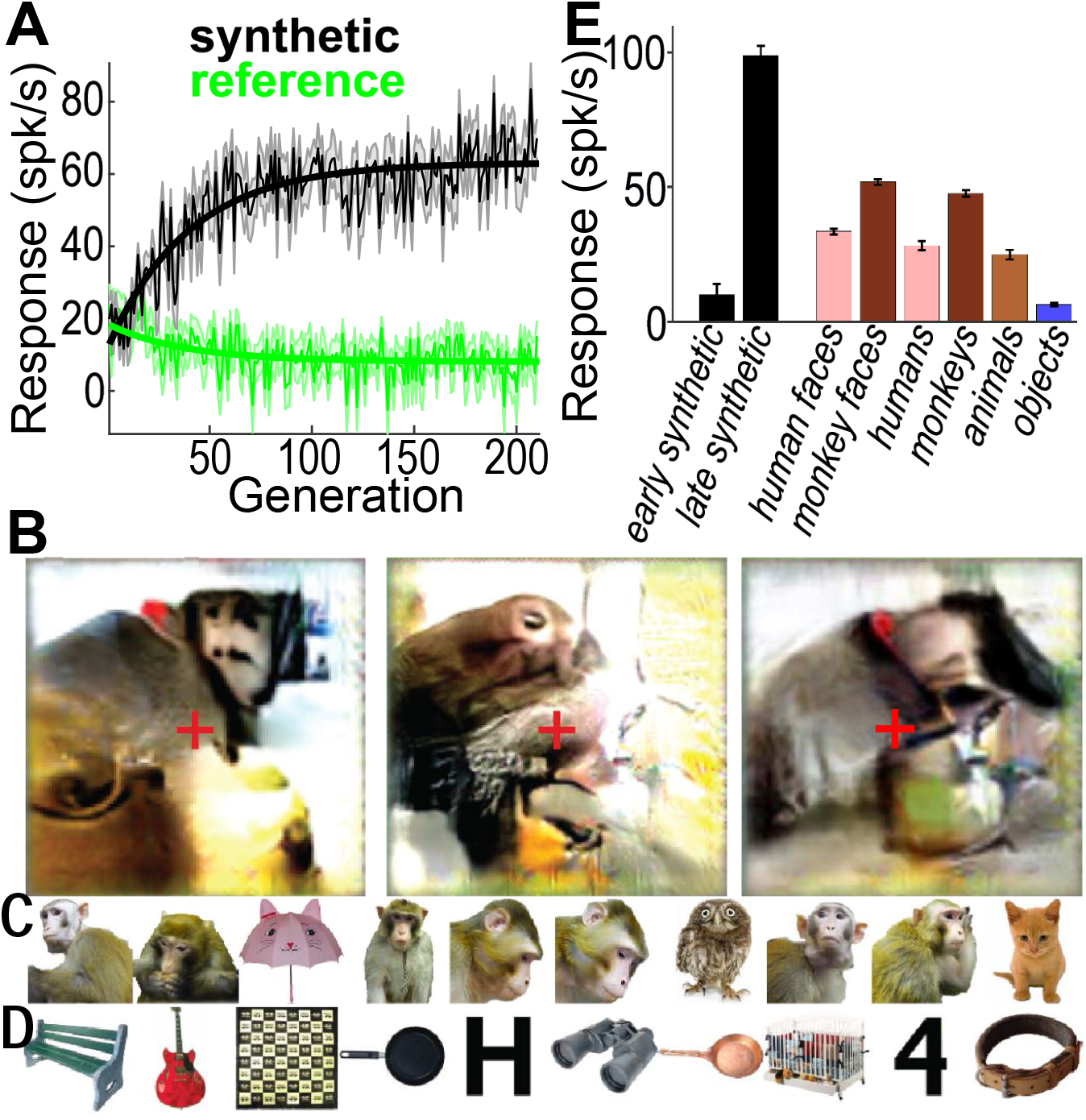
Evolution of synthetic images by maximizing responses of single neuron Ri-10 (same unit as Figure 3) (**A**) Mean response to synthetic (black) and reference (green) images for every generation (spikes per s + sem). Solid straight lines show an exponential fit to the response change over the experiment. (**B**) Last-generation images evolved during three independent evolution experiments; the leftmost image corresponds to the evolution in (**A**); the other two evolutions were carried out on the same single unit on different days. Red crosses indicate fixation. The left half of each image corresponds to the contralateral visual field for this recording site. Each image shown here is the average of the top 5 images from the final generation. (**C&D**) Selectivity of this neuron to 2,550 natural images. (**C**) Top 10 images from this image set for this neuron. (**D**) Worst 10 images from this image set for this neuron. The entire rank ordered natural image set is shown in Figure S1. (**E**) Selectivity of this neuron to different image categories (mean+sem). The entire image set comprised 2,550 natural images plus the best synthetic image from each of the 210 generations; 10–12 repetitions each; early synthetic was the first 10 generations and late the last 10.

We conducted independent evolution experiments with the same single unit on different days, and all final-generation synthetic images featured a brown object against a uniform background, topped by a smaller round pink/brown region containing several small dark spots; the object was centered toward the left half of the image, consistent with the recording site being in the right hemisphere (Figure 4B). The synthetic images generated on different days were similar by eye, but not identical, presumably due to response variability and/or stochastic paths explored by the algorithm in the neuron’s response landscape. Regardless, given that this unit was located in PIT, just anterior to the tip of the inferior occipital sulcus and thus early in the visual hierarchy, it was impressive that it reproducibly evolved images that contained such complex motifs and that evoked such high firing rates. Two days following the evolution experiment in Figure 2 this unit was screened with 2,550 natural images, including animals, bodies, food, faces and line drawings, plus the top synthetic images from each generation. Among the natural images this neuron responded best to monkeys and monkey faces (Figure 4C). Of the 10 individual natural images in this set giving the largest responses, five were of the head and torso of a monkey. The worst natural images were of inanimate and rectilinear objects (Figure 4D). The strongest stimuli for this neuron, by far, were the late-generation synthetic images (Figure 4E). Figure 5A shows a histogram of response magnitudes for this same PIT cell to the top synthetic image in each of the 210 generations and responses to each of the 2550 natural images (collected 2 days later). Early generations are indicated by lighter gray and later by darker, so it is apparent that later generation synthetic images give larger responses. Figure 5B shows the same kind of histogram for an experiment on a cell in CIT of monkey Ge.

**Figure 5.**
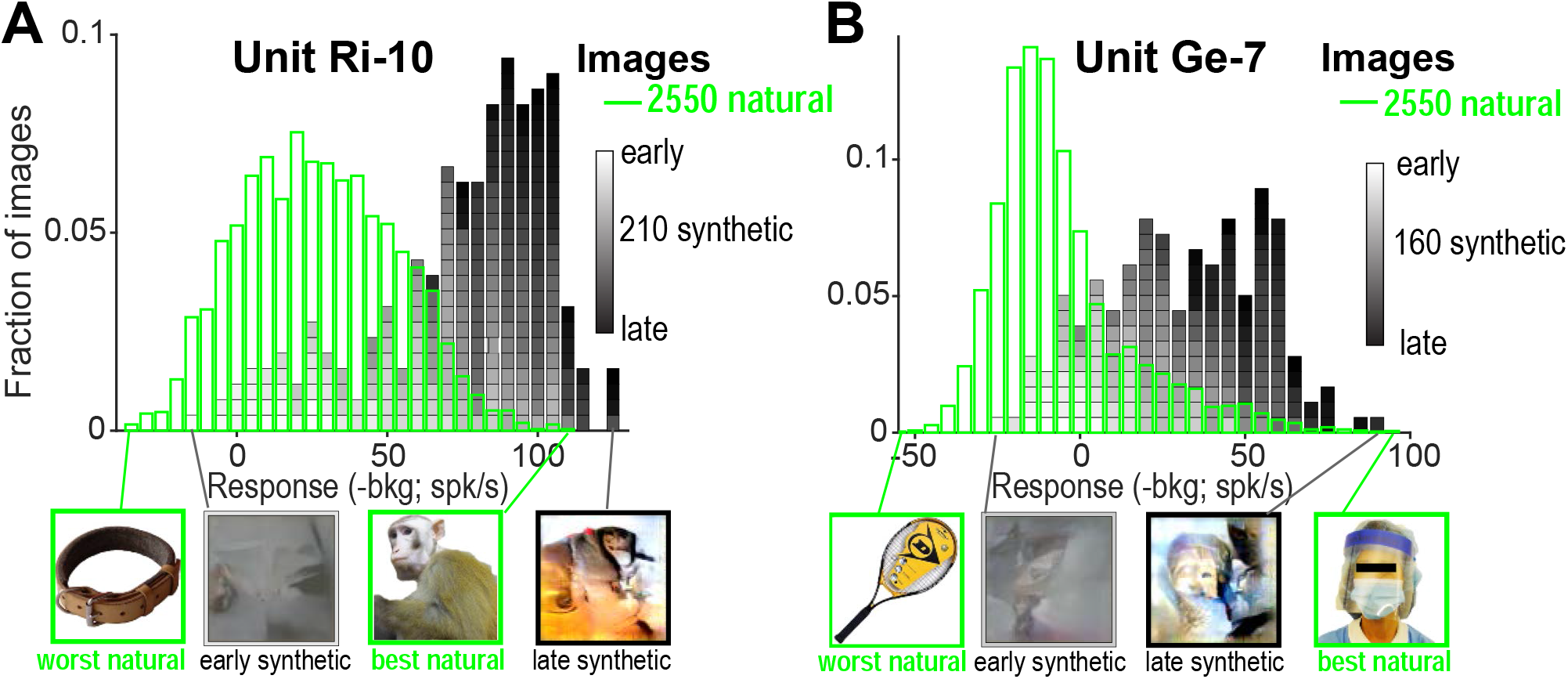
Histogram of response magnitudes to natural (green) and synthetic (gray-to-black) images for evolutions for two cells. (**A**) Histogram for unit Ri-10 (same unit as Figures 3 & 4). Below the histogram are shown the best and worst natural and synthetic images. (**B**) Same for unit Ge-7.

### 2.3. Evolution of synthetic images in other neurons

We conducted 35 independent evolution experiments on single- and multi-unit sites in IT in five different monkeys. IT neurons guided the evolution of images that varied from experiment to experiment, but retained consistent features for any given recording site, features that reflected each neuron’s selectivity to natural-image sets. Figure 6A shows the final-generation synthetic images from two independent evolution experiments for one IT site in each of four monkeys, along with each site’s top 10 natural images (for monkeys Gu and Y1 these were from the 108 reference images shown interleaved with the synthetic images during the evolution experiment; for monkeys Ri and Ge these were from the 2550 set of natural images shown in an independent experiment). In each case a reproducible figure emerged in the left half of the synthetic images, corresponding to the contralateral visual field. In three monkeys (Ri, Gu, and Ge) screening with natural images indicated that the arrays were located in face-preferring regions, but in the fourth animal (Y1), the array was in a place-preferring region (Figure 6B). During almost all the evolutions, the synthetic images evolved gradually to become very effective stimuli. To quantify the change in stimulus effectiveness over each experiment we fit an exponential function to the mean firing rate per generation (as in Figure 3A), separately for synthetic and reference images. Synthetic-image rates changed between 25.5 and 81.4 spikes per s per animal (Figure 6C, colors); individual amplitude values were significantly different from zero in 33 out of 35 experiments (95% CI of amplitude estimate not including zero per bootstrap test). In contrast, the same neurons showed stable or slightly decreasing responses to the reference images across generations, (amplitudes estimates per array population ranged from −10.4 to 8.7, significant in 17/35 experiments, Figure 6C, Table S1). Thus IT neurons could consistently guide the evolution of strongly effective images, despite minor adaptation. Moreover, as we show below, these synthetic images were often more powerful stimuli for these neurons than were the best natural images we could find, despite the fact that the synthetic images were far from naturalistic.

**Figure 6.**
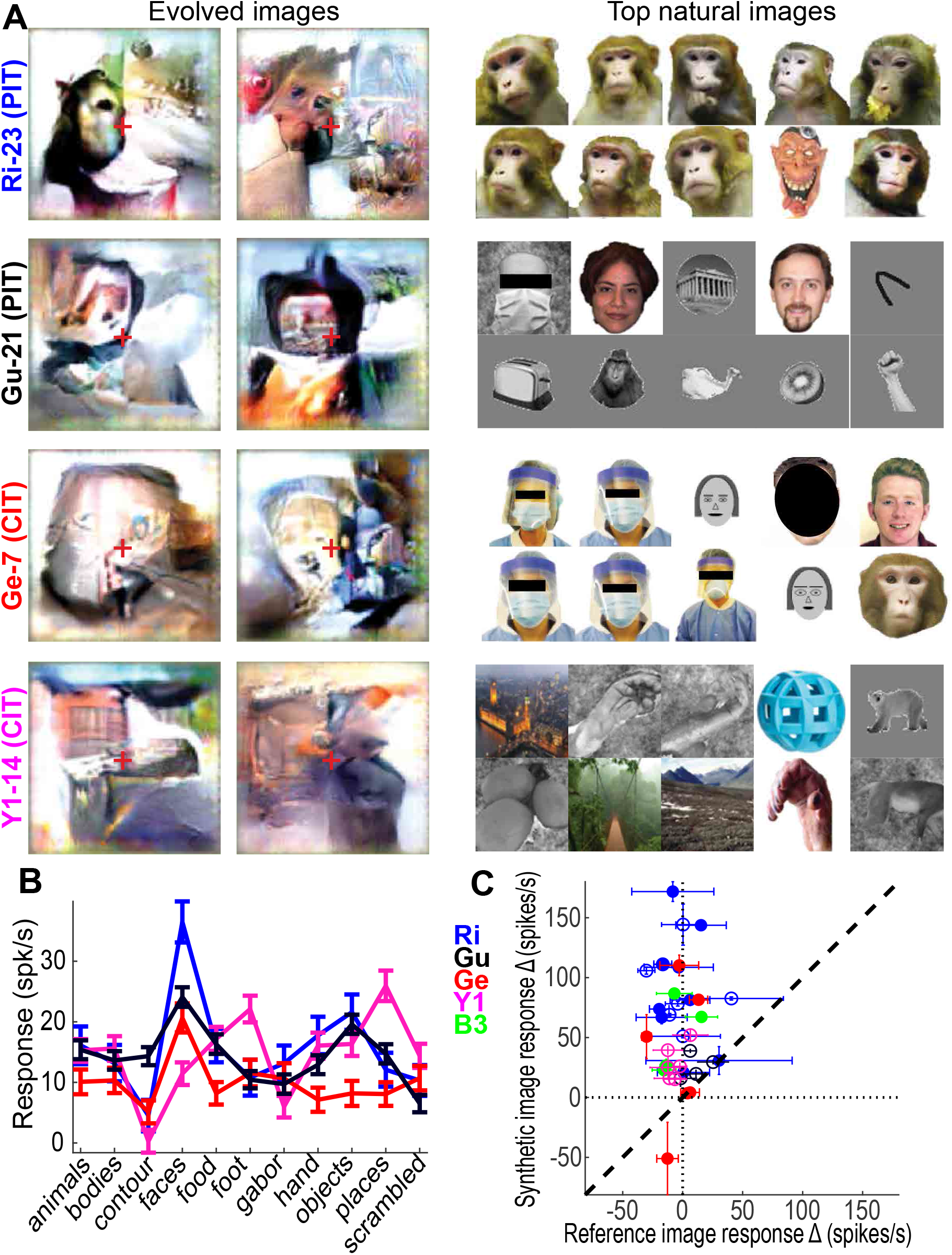
Evolution of synthetic images by maximizing firing rate; other IT neurons. (**A**) Last-generation synthetic images from two independent evolution experiments for a single chronic recording site in the right hemisphere of each of 4 different animals. To the right of the synthetic images are shown the top 10 images for each neuron from a natural image set. Red crosses indicate fixation. (**B**) Category selectivity of the arrays in (**A**) (mean + sem; no data for B3). (**C**) Responses to synthetic vs reference images over generations. Each point shows the slope of a linear regression of spikes per s per generation for reference (x axis) and synthetic images (y axis) for an individual experiment (±sem). Solid circles indicate single units; open circles multi-units.

### 2.4. Predicting neuronal responses to a novel image from its feature similarity to the evolved stimuli

If these evolved images are telling us something important about the tuning properties of IT neurons, then we should be able to use them to predict neurons’ responses to novel images. The deep generator network had been trained to synthesize images from their encoding in layer fc6 of AlexNet (4096 units), so we used the fc6 space to find natural images similar to the synthetic images. This would allow us to search for effective natural images in a data set much larger than we could possibly screen by showing hand-picked images to neurons. In particular, we could ask whether a neuron’s response to a novel image was predicted by the distance in fc6 space between the novel image and the neuron’s evolved synthetic image. To do this we calculated the activation vectors of the evolved synthetic images in AlexNet fc6 and searched in image space for images with similar fc6 activation vectors. We used 2 databases for this: the first comprised ~60,000 images collected in our laboratory over several years, and the second set comprised 100,300 randomly sampled images from ImageNet 2012 (https://arxiv.org/abs/1409.0575; we included 100 images from each of its 1000 categories, in addition to the ImageNet categories of *faces* [ID *n09618957*], *macaques* [ID *n02487547*], and 100 images of local animal care personnel wearing personal protective equipment). In experiments using the latter database, there were 100,300 activation vectors which were ranked by their proximity to the activation vector of each synthetic image.

First, we focus on the evolution experiment for PIT single unit Ri-17. This cell evolved a discrete shape near the left top of the image frame, comprising a darkly outlined pink convex shape with two dark circles and a dark vertical line between them (Figure S2A). When tested with a 2,550 natural image set, this neuron responded best to images of monkeys, dogs, and humans (Figure S2B). We propagated this evolved image through AlexNet along with the 100,300 ImageNet examples, and ranked all the fc6 vectors by their Pearson correlation to the evolved image vector. We identified the closest, middle and farthest 100 matches. The synthetic image showed a mean vector correlation of 0.38 to the closest images, 0.06 to the middle and −0.14 to the farthest images. The 9 nearest ImageNet images were cats, dogs and monkeys (Figure S2C). To visualize the common shape motifs of this image cluster, we identified the individual fc6 units most strongly activated by the synthetic image and used activation maximization (*deepDreamImage.m; Matlab, Natick, MA*) to generate examples of those fc6 units’ preferred shapes. All the units were filters for round tan/pink regions with small dark spots (Figure S2D). To rule out that these matches could be due to an overrepresentation of animals in ImageNet, we also looked at the least correlated matches, which were indeed not animals, but were pictures of places, rectilinear textures, or objects with long straight contours (Figure S2E).

We applied this image-search approach to all evolution experiments by identifying the top 100 matches to every synthetic image in fc6 space (the Pearson correlation coefficients of these images ranged from 0.30 to 0.61, median 0.36) and visualized the WordNet (Russakovsky et al., 2015) labels of the matching images via word clouds. In monkey Ri, whose array showed classical preferences for faces, the categories that best matched the synthetic images were “macaques”, “toy terrier” and “Windsor tie” (the latter containing images of faces and bodies) (Figure S2F); in contrast, in monkey Y1, where most of the neurons in the array had shown classical preferences for places, the categories that best matched were “espresso maker,” “rock beauty” (a type of fish) and “whiskey jug” — not an informative set, but by inspection these images all contained extended contours (Figure S2G). We confirmed this trend that WordNet categories matched evolved images by quantifying the WordNet hierarchy labels associated with every matched natural image (Table S2).

To find out whether distance in fc6 space between a novel natural image and a neuron’s evolved synthetic image predicted that neuron’s response to that novel image, we first performed 3–4 independent evolution experiments using the same (single- or multi-) unit in each of three animals. After each evolution, we took the top synthetic image from the final generation and identified the top 10 nearest images in fc6 space, 10 images from the middle of the distance distribution and the farthest 10 (most anti-correlated images) (Figure 7A). We then presented these images to the same IT neurons and measured the responses to each group (*near, middle, far*) as well as to all 40 evolved images of the last generation. Figure 7B shows that synthetic images gave the highest responses, the nearest natural images the next highest responses, and the middle and farthest images the lowest. Thus distance from the evolved synthetic image in fc6 space predicted responses to novel natural images. But importantly, responses to the synthetic images were the highest of all. Consistent with this, in the evolution experiments we interleaved natural images along with synthetic images, and, in general, late-generation synthetic images elicited higher firing rates than did the natural images. When comparing the cells’ maximum responses to natural vs. synthetic images, cells showed significant differences in 16 of 35 experiments (P < 0.03, permutation test after false discovery correction), and, in all but two cases, synthetic images evoked the maximum response. The single interesting exception was when we compared synthetic images evolved by multi-unit site 7 in monkey Ge (illustrated in Figure 5B and in the 3rd row of Figure 6A) against 2,550 natural images, two days after the original evolution experiment. This same site responded slightly better (by an average of four spikes per s, or 3.7% of its maximum rate) to images of a familiar animal-care person wearing the protective mask and gown she typically wears in monkey rooms, so this image represents something the monkey experiences frequently. Even in this case, one clear advantage of the evolved image is that it revealed the specific shape patterns within the natural image that were diagnostic of neuronal selectivity, stripped of incidental information. See Table S3 for further quantification of natural and synthetic image responses.

**Figure 7.**
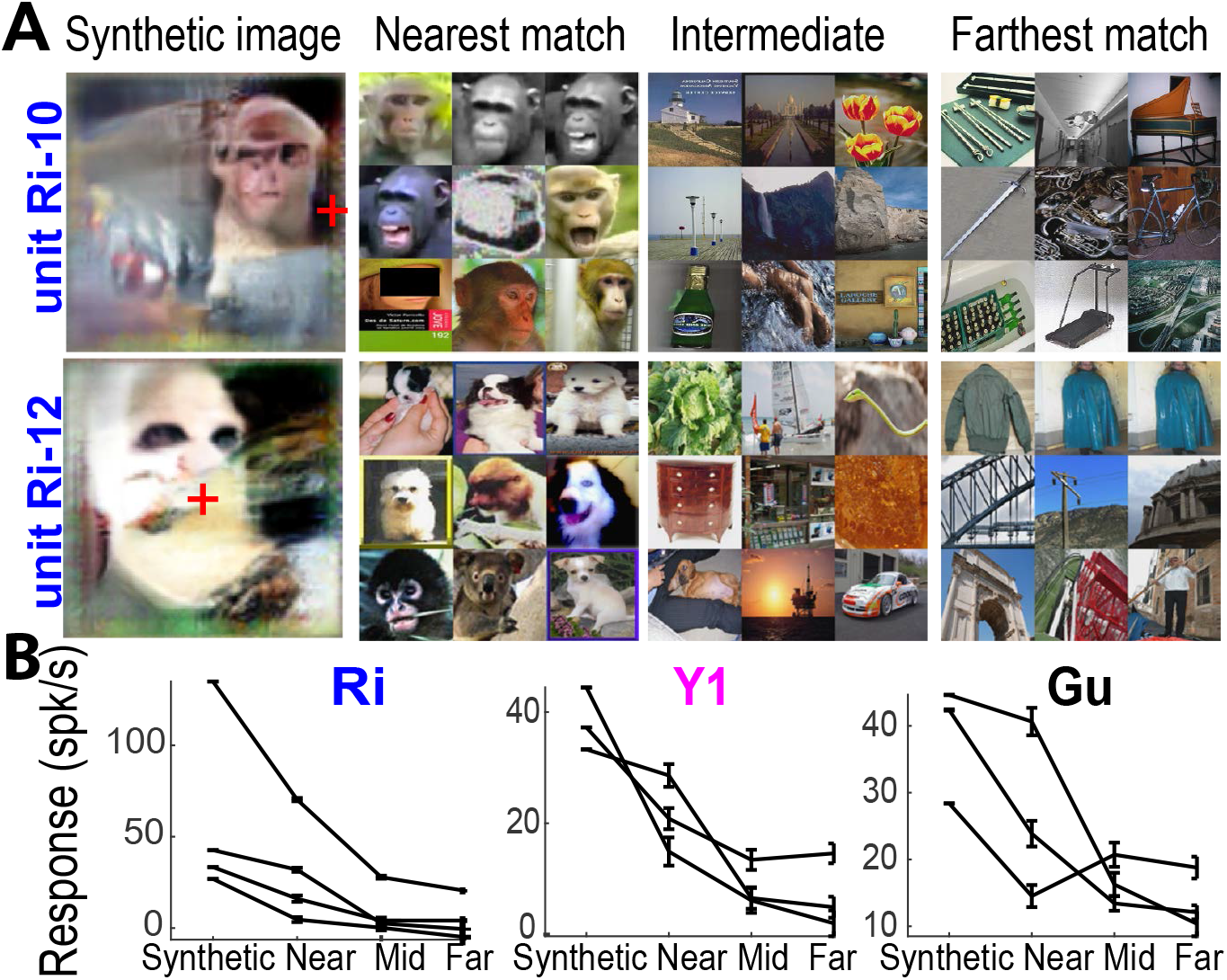
Using AlexNet to predict responses to novel images. (**A**) Final-generation synthetic images from units Ri-10 and Ri-12 and the closest, intermediate, and farthest 9 images from the image set for each. For unit Ri-10 we used the 60,000 image database, and for unit Ri-12 the 100,300 image database. (**B**) Responses from each unit to the last-generation evolved synthetic images compared to the nearest, intermediate, and farthest images in the fc6 space of ImageNet (mean + sem).

### 2.5. Invariance to evolved vs natural images

IT neurons retain selectivity despite changes in position, size and rotation (Ito et al., 1995; Kobatake & Tanaka, 1994), although it has been reported that more selective neurons are less transformation-invariant (Zoccolan et al., 2007). To compare the invariance of IT neurons to synthetic and natural images, we presented 3 natural and 3 evolved synthetic images at different positions, sizes and fronto-parallel rotations in two animals (monkeys Ri and Gu). The natural images were chosen from the nearest, middle, and farthest matches from ImageNet. The synthetic images were chosen from the final generation. Every image was presented at three positions relative to the fovea: (−2.0°, −2.0°), (−2.0°, 2.0°) and (0.0°, 0.0°); three sizes (width of 1°, 2°, 4°) and 4 rotations (0°, 22°, 45° and 80°, counterclockwise from horizontal) (Figure S3A). Invariance was defined as the similarity (Pearson correlation coefficient) in the neuron’s rank order of preferences for different images under different transformation conditions (the more similar the rank order, the higher the invariance). The rank order was better maintained across transformations for the natural images than for the synthetic images (Figure S3B). Thus the degree of invariance for these neurons changed depending on the stimulus set, and the neurons were the least invariant for the more optimal synthetic images. This result suggests that the degree of invariance measured for particular neurons may not be a fixed feature of that neuron, but rather may depend on the effectiveness of the stimulus used to test the invariance.

### 2.6. Generative evolutions from populations of neurons

Single- and multi-units in IT successfully guided the evolution of synthetic images that were stronger stimuli for the neuron guiding the evolution than large numbers of natural images. Each of the chronically implanted arrays had up to 32 visually responsive sites, spaced 400 μm apart. To see if our technique could be used to characterize more coarsely sampled neuronal activity than a single site we asked whether we could use all 32 sites on an array. We conducted a series of evolution experiments in 3 monkeys (Ri, Gu, and Y1) guided by the average population response across the array. Evolution experiments for all 3 monkeys showed increasing responses to synthetic images over generations compared to reference images: The median population response change to synthetic images for monkeys Ri, Gu and Y1 were 0.04 spikes per s per generation (0.01–0.10, 25th–75th percentile), 0.17 spikes per s per generation (0.08–0.31, 25th–75th %tile) and 0.19 spikes per s per generation (0.09–0.25, 25th–75th %tile). In these population-based evolutions, between 33.7% to 88.8% of individual sites within the population per animal showed increases in firing rate statistically different from 0 (*P* < 0.01 Student’s t-test after false discovery correction). Therefore larger populations of IT neurons could successfully create images that were on average strong stimuli for that population. When the populations were correlated in their classical preferences, the synthetic images were consistent with those evolved by individual single sites in the array: for example, in monkey Ri, the population-evolved images contained shape motifs commonly found in ImageNet pictures labeled “macaques,” “wire-haired fox terrier,” and “Walker hound.” This suggests that the technique could be used with coarser sampling techniques than single-unit recordings, such as local field potentials, electrocorticography electrodes, or even functional magnetic resonance imaging.

### 2.7. Testing the generative evolution algorithm using the ground truth of primary visual cortex

We recorded from one single unit and three multiunit sites (six evolution experiments total) in monkey Vi, which had a chronic microelectrode array in V1. The stimuli were centered on each receptive field (measuring 0.79° square root of the area), but were kept at the same size as in the IT experiments (3°x3°-wide). In addition to the synthetic images, we interleaved reference images of gratings (3°x3° area) of different orientations (0°, 45°, 90° and 135°) and 2–3 spatial frequencies (~0.5, 1 and 2 cycles per °) at 100% contrast. In all experiments, neurons showed an increase in firing rate over generations to the synthetic images (median 84.0 spikes per s per gen (77.4–91.2, 25th–75th percentile; Table S1). Thus on average, V1 sites, like IT, responded best to late-generation synthetic images (Table S3). To measure the distribution of orientations of the region of the synthetic images that fell within each V1 receptive field (~0.8°×0.8°), we performed a discrete Fourier transform analysis on the central 0.8°×0.8° of the synthetic images and correlated the resulting spectrogram to the spectrograms expected from 16 gratings with orientations ranging from 0° to 135°. Across experiments, the mean correlation between the orientation content profile of the patch and the orientation tuning measured from the gratings was 0.59±0.09 (mean ± SEM), compared to 0.01±0.26 for a shuffled distribution (*P* values ≤ 0.006 in 5/6 experiments, permutation test, *N_iterations_* = 999). Thus V1 neurons indeed guided the evolution of images to generate synthetic textures that were dominated by the neurons’ independently measured preferred orientation.

## 3. Discussion

We introduce a new algorithm for studying the response properties of visual neurons using a vast and unbiased generative image space. The algorithm starts with random shapes and evolves those shapes based on neuronal responses. The generative algorithm led to images that elicited large responses in V1, in different parts of IT, in single-units, in multi-units, and in average population responses. Remarkably, the generative network evolved stimuli that evoked higher responses than the best natural images found by an extensive exploration of large image sets. In some instances, the evolved stimuli contained features of animal faces, bodies, and even depictions of animal-care staff known to the monkeys, consistent with theories of tuning to object categories and faces. However, often the evolved stimuli were not of identifiable objects or object parts, suggesting that current theories of visual cortex have missed aspects of neuronal response properties.

It is unclear whether this approach has uncovered globally optimum stimuli, if such things exist, for these cells. It is not clear there can be a single global optimum stimulus for a particular neuron. Different evolutions for the same units yielded synthetic images that looked different by eye (*e.g*. Figure 4B), as would be expected from a cell that shows invariance to nuisance transformations like position, color, or scale. Complex cells even in early visual areas respond equally well to the same stimulus in multiple locations within a cell’s receptive field (Hubel & Livingstone, 1985; Hubel & Wiesel, 1968), and this complexification/OR gate-like operation is likely to occur at multiple levels in the hierarchy (Hubel & Livingstone, 1985; Riesenhuber & Poggio, 1999), so it follows that multiple different feature combinations could strongly activate a particular neuron in IT. What are these powerful stimuli? To start to answer this, we looked for images in AlexNet layer fc6 space that were nearest to the evolved images, because the network had been trained on that space and high layers of this network are correlated with perception (Yamins et al., 2014). Distance in fc6 space predicted neuronal selectivity. For neurons classically defined as monkey- and face-preferring (Desimone et al., 1984; Perrett et al., 1982), the closest images to the evolved ones were macaques and dogs or contained faces and torsos of monkeys and other mammals. In contrast, for place-preferring neurons, the nearest images instead included a variety of objects including espresso makers and moving vans. Although the evolved synthetic images were not life-like, or sometimes even identifiable objects, they nevertheless tell us something quite novel about what information might be encoded by IT neurons. PIT neurons have been reported to be selective for low-level features like radial gratings (Pigarev et al., 2002), color (Zeki, 1977), or single eyes (Issa & DiCarlo, 2012). Our experiments reveal a more intriguing view since both PIT and CIT neurons evolved complex synthetic images. The unrealistic nature of the evolved images, plus the fact that these images were more effective than any images in an extensive natural-image database, suggest that we may have evolved “super-stimuli” that are more effective for a particular neuron than anything the animal would ever encounter. This is consistent with the idea that neurons are tuned to abstract parameters, such as axes (Chang & Tsao, 2017; Freiwald et al., 2009; Leopold et al., 2006) that distinguish between things in the environment, rather than being tuned to the things themselves. That is, neuronal responses may carry information about an object in the environment, not a veridical representation of it. These results complement classical methods for defining neuronal selectivities and provide the potential for uncovering internal representations in any modality that can be captured by generative models.

## Acknowledgment

This work was supported by NIH grants R01EY16187, R01EY25670, R01EY011379, P30EY012196, R01EY026025, NSF STC award CCF-1231216 to CBMM at MIT

## Author Contributions

GK and WX conceived of the approach, WX implemented the network algorithm, CP & PS adapted the algorithm to neurophysiology, PS, ML, TH & CP collected and analyzed data, CP & ML wrote the paper, and ML acquired funding.

## Supplementary Materials

METHODS Figures S1-S3 Tables S1-S4

## 4. Methods

### 4.1. Deep Generative Neural Network

The pre-trained generative network was downloaded from the authors’ website (lmb.informatik.uni–freiburg.de/resources/software.php) and used without further training with the Caffe library (caffe.berkeleyvision.org) in Python. To synthesize an image from an input image code, we forward propagated the code through the pre-trained generative network and clamped the output image pixel values between 0 and 1. Some images synthesized by the network contained a patch containing a stereotypical shape that occurred in the center right of the image. This was identified as an artifact of the network commonly known as “mode collapse” and it appeared in the same position in a variety of contexts, including a subset of simulated evolutions. This artifact was easily identifiable and it did not affect our interpretations. In the future, more modern GNN training methods can avoid this problem (personal correspondence with Alexey Dosovitskiy).

### 4.2. Initial generation

The initial generation of image codes was constructed from a set of Portilla and Simoncelli textures, derived from randomly sampled photographs of natural objects in a gray background. We started from all-zero codes and optimized for pixelwise loss between the synthesized images and the target images using backpropagation through the network for 125 iterations, with a learning rate linearly decreasing from 8 to 1 ∗ 10^−10^. The resulting image codes produced blurred versions of the target images, which was expected from the pixelwise loss function and accepted because the initial images were intended to be quasi-random textures. We also tried initializing the image codes with the AlexNet fc6 encoding of the target initial images, but the resulting evolutions were qualitatively similar.

### 4.3. Genetic algorithm

The function began with an initial population of 40 image codes (‘individuals’), each consisting of a 4096-dimensional vector (‘genes’) and associated with a synthesized image. Images were presented to the subject, and the corresponding spiking response was used to calculate the “fitness” of the image codes by transforming the firing rate into a Z-score within the generation, scaling it by a selectiveness factor of 0.5, and passing it through a softmax function to become a probability. The 10 highest-fitness individuals were conserved to the next generation without recombination or mutation. Another 30 children image codes were produced from recombinations between two parent image codes from the current generation, with the probability for each image code to be a parent being its fitness. The two parents contributed unevenly (75%:25%) to any one child. Individual children genes had a 0.25 probability of being mutated, with mutations drawn from a 0-centered gaussian with standard deviation 0.75. Hyperparameter values were not extensively optimized. All source code is available per request.

### 4.4. Generative evolution using CaffeNet

We selected 100 random units each in 4 layers in CaffeNet as targets for evolution. For convolutional layers, only the center unit in each channel was considered. Each unit was evolved for 500 generations, 10,000 image presentations total. The best image in the last generation was used in the analysis, although most of the total activation increase was achieved by 200 generations. As a control, we recorded activations of the units to all 1431167 images in the ILSVRC2012 dataset (Deng et al., 2009), including the training set of CaffeNet. To visualize the ground truth best in CaffeNet layer conv1, 11×11×3 images were produced from the 11×11×3 filter weights according to *image*(*x, y, c*) = 0.5 + *sign*(*weight*(*x, y, c*))/2, because this is a linear transformation. In other words, positive weights corresponded to a pixel value of 1 and negative weights 0. To further visualize the magnitude of the weights, each pixel (x,y) was made transparent to a grey checkerboard in inverse proportion to its contribution to the overall activation.

### 4.5. Neurophysiology

All procedures were approved by the Harvard Medical School Institutional Animal Care and Use Committee, and conformed to NIH guidelines provided in the Guide for the Care and Use of Laboratory Animals. **Visual stimuli:** We used MonkeyLogic2 as the experimental control software. Images were presented on an LCD monitor 53 cm in front of the monkey at a rate of 100 ms on, 100–200 ms off. **Behavior:** Six adult male macaques (9–13 kg) were trained to perform a fixation task. They fixated on a 0.2°-wide fixation spot in the middle of the screen. Eye position was monitored using an ISCAN system (Woburn, MA). Animals were rewarded with a drop of water or juice for maintaining fixation within 2.0° of the fixation spot for 2–7 image presentations; the interval was gradually decreased over the experimental session as the monkey’s motivation decreased. **Recording arrays:** Monkeys Ri, Gu, Ge, and Y1 were implanted with custom floating microelectrode arrays manufactured by MicroProbes for Life Sciences (Gaithersburg, MD); each had 32 platinum/iridium electrodes per ceramic base, electrode lengths of 2–5 mm, impedances between 0.7–1.0 MΩ. Monkey V1 was implanted with a 128-channel Utah array (Blackrock microsystems, Salt Lake City, Utah). Monkey B3 had an acute recording chamber, and neuronal activity was recorded using a 32 channel NeuroNexus Vector array (Ann Arbor, Michigan) that was inserted each recording day. Neural signals were amplified and extracellular action potentials were isolated using the box method of an online spike sorting system (Plexon, Dallas, TX). Spikes were sampled at 40 kHz. **Surgical procedures:** All animals were implanted with custom-made titanium or plastic headposts before fixation training. After several weeks of fixation training, the animals underwent a second surgery for array implantation. PIT insertion sites were just anterior to the inferior occipital sulcus; CIT sites were on the lower lip of the STS 6–8 mm anterior to the interaural line. All surgeries were done under full surgical anesthesia using sterile technique.

### 4.6. Data analysis

**Quantification of spike rate changes during evolution experiments.** We defined the neuronal response as the spike rate measured in the 70–200 ms time window after image onset and subtracted the rate in the 0–80 ms window. In the evolution experiments, there were 40 synthetic and 40 reference images per generation, each presented once. Reference images are from various sources, including personal photos, Imagenet (Deng et al., 2009), Caltech-256 (Griffin et al., 2007), and PICS (http://pics.stir.ac.uk/). To track firing rate change per generation, we averaged responses to all 40 synthetic and 40 reference images separately. We used two statistical measures to estimate the inter-generational changes in response, fitting the mean response/generation curve separately with a 1) linear regression function and 2) a decaying exponential function −*a e*^(−*x*/*τ*^) + *c*. The exponential decay description start at the first block with rate c and asymptotically approach the rate a+c (a being the amplitude increase or decrease). Tau controls its slope. We restricted the amplitude change to be within physiologically viable values (±1.0x absolute maximum difference of any generations’ rate for this day). To assess statistical significance, we generated new mean rate per generation curves by resampling responses (with replacement) from each of the 40 synthetic and 40 reference image presentations within one generation, then fit the exponential function each time (N = 500 repetitions). The 95% confidence intervals reported are the 0.025/500 and 0.975/500 (12th and 488th) value of the bootstrapped distribution. We used responses from all generations except for one experiment in monkey B3, where single-unit isolation instability restricted interpretable data to generations 15–70. **Quantification of natural image frequency labels to the evolved images.** Every evolved image was propagated into AlexNet and its fc6-layer activation was compared to those of 100,300 natural images sampled from ImageNet (Deng et al., 2009). Every photograph in ImageNet is labeled by categories defined by WordNet, a hierarchically organized lexical database (7). After ranking every natural image by its proximity to the evolved image (via Pearson correlation coefficient), we used a tree search algorithm to crawl through each labeled image’s label hierarchy for the specific search terms “macaque,” “monkey”, “face,” and “appliance” (“place” was not used because place images often contained people). We measured the frequency of labels associated with all evolved images for every subject. To estimate confidence intervals for every observed frequency, we re-sampled the top matches to each evolved image 200 times (with replacement) and repeated the analysis. To test if the frequency of photographs labeled “monkeys” and “appliance” were statistically different between subjects Ri and Y1, we used a permutation test. The null hypothesis was that these frequency values arose from the same distribution, so we shuffled labels from the Ri and Y1 populations, sampling twice with replacement, and measured the difference, 500 times. We then compared the observed difference in frequency values with the distribution.

## A. Supplementary Materials

**Figure S1.**
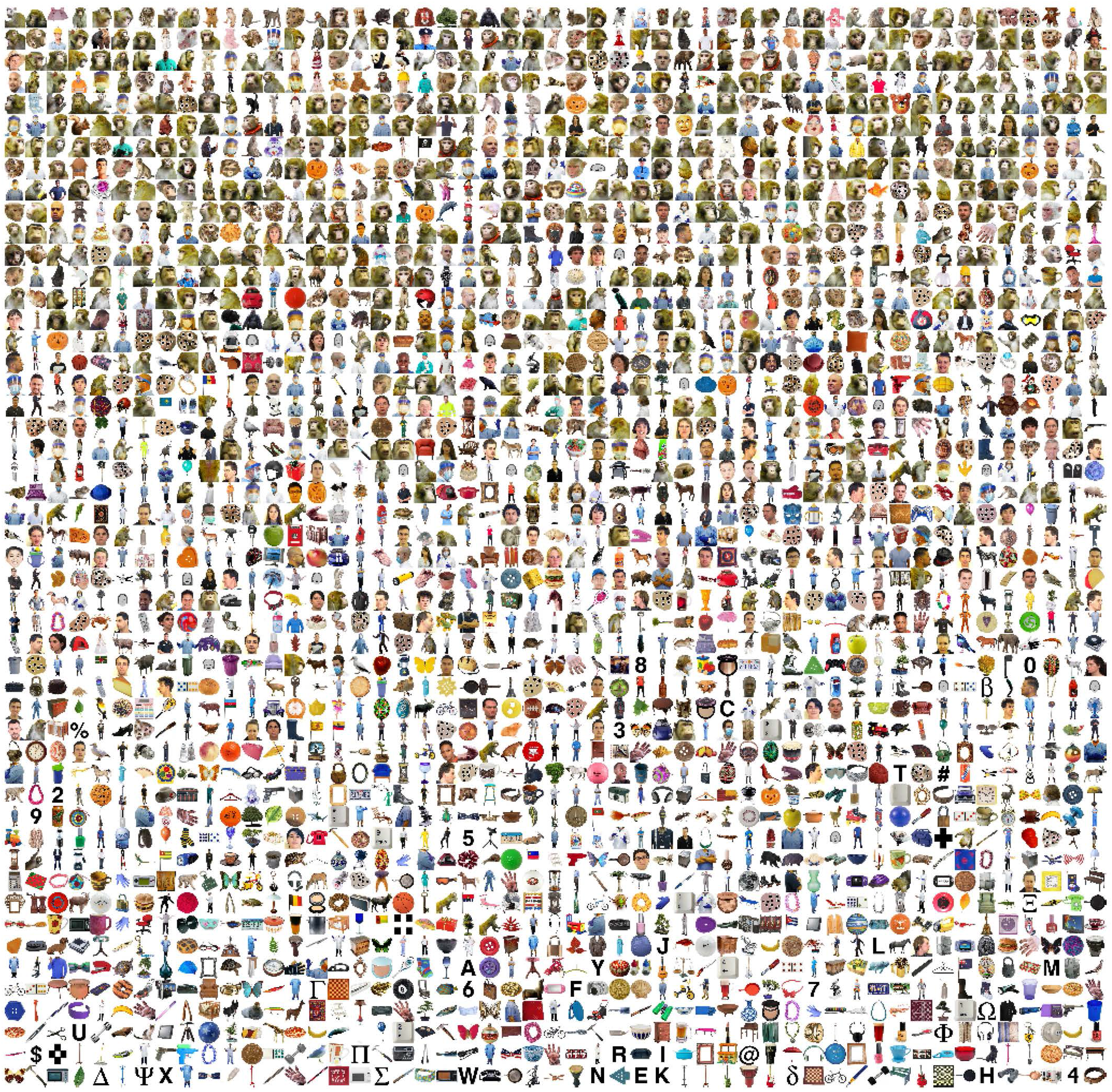
Rank order from highest (top left) to lowest (bottom right) responses of unit Ri-10 to 2550 images. Responses were the number of spikes between 70 and 250 ms after stimulus onset, minus baseline (average number of spikes from 1–50 ms after stimulus onset). The average response to these images by category is shown in Figure 3E.

**Figure S2.**
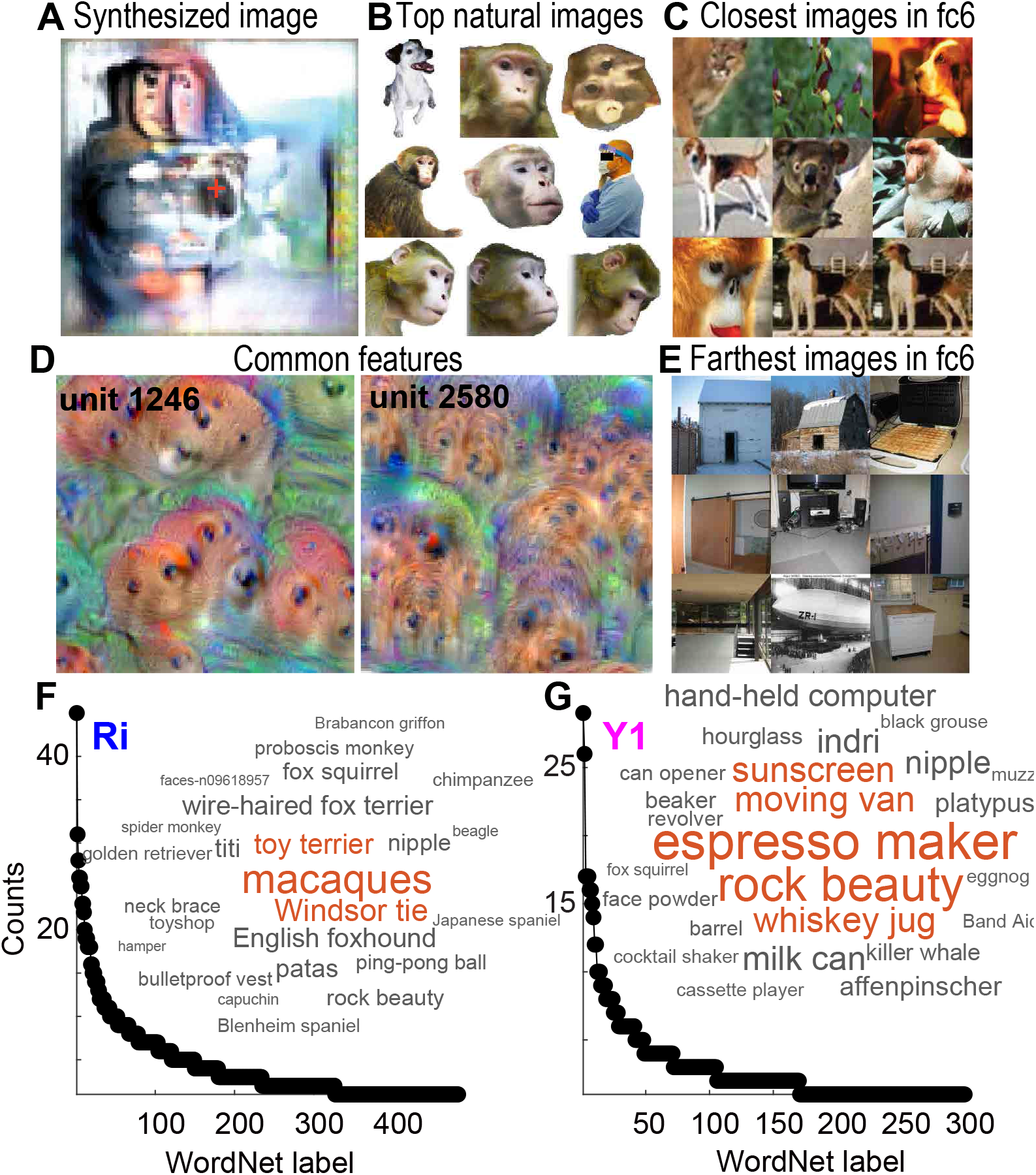
Exploration of image space of synthetic images. (**A**) Example image evolved by PIT single unit Ri-17. Its receptive field encompassed the contralateral (left) side of the image; red cross indicates fixation. (**B**) Top 9 images from our 2550 image set for this neuron. (**C**) Closest ImageNet picture matches based on AlexNet fc6 distance. (**D**) Common shape features in (C), as encoded by fc6 units that showed highest activations in response to the synthesized image. (**E**) Furthest ImageNet picture matches. (**F**) word cloud and histogram showing counts of ImageNet labels of the top 150 matches for all experiments for all 14 visually responsive sites in the array in monkey Ri. (**G**) Same, but for images evolved by monkey Y1 recording sites, which preferred images of places.

**Figure S3.**
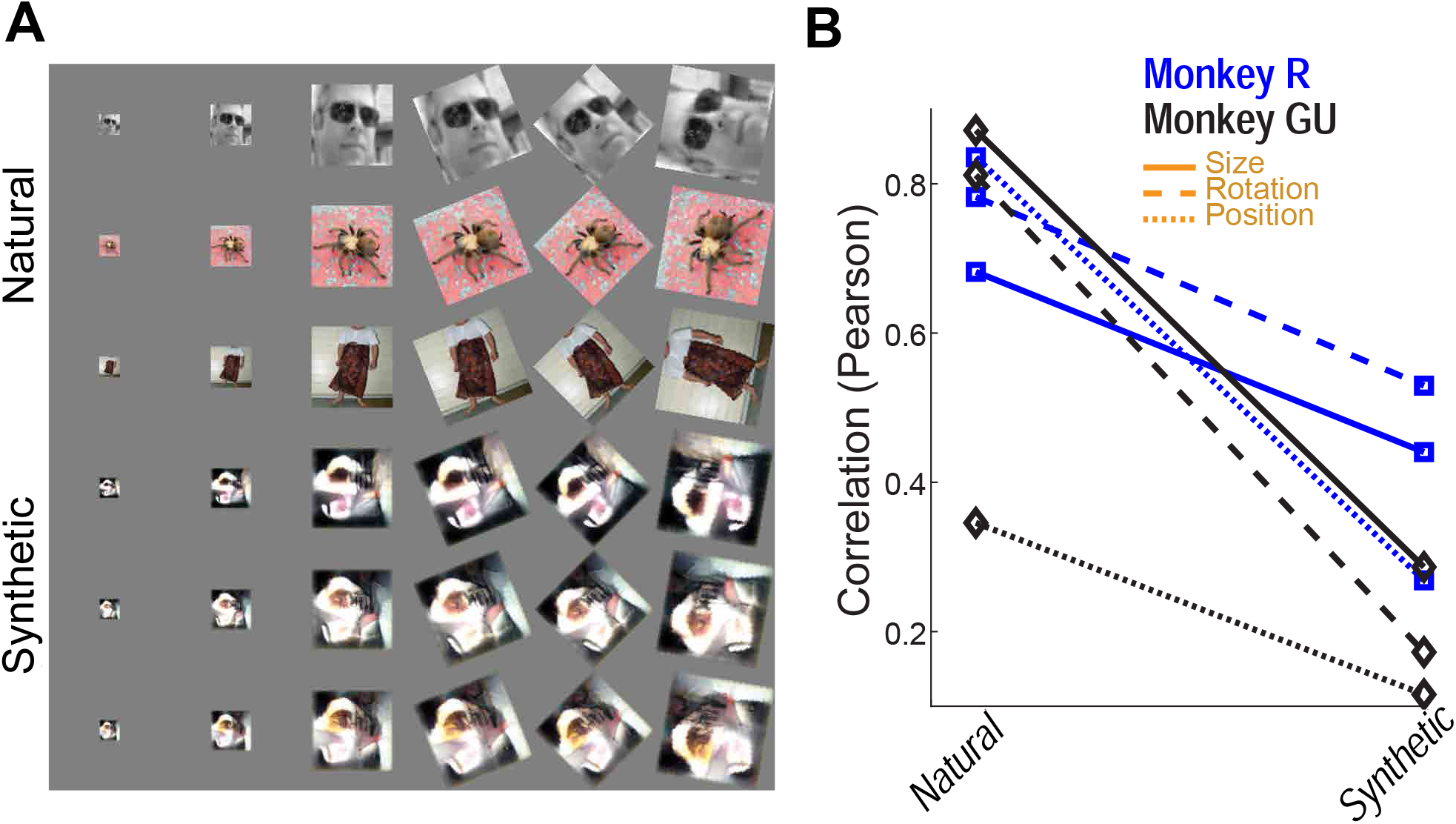
Size, rotation, and position invariance. (**A**) Transformations applied to natural images and to synthetic images evolved by PIT units (the natural images were nearest, intermediate, and farthest fc6 matches to the evolved image). Images varied in size, rotation and position. (**B**) Correlation of rank order preferences for natural vs. synthetic images across transformations and monkeys.

**Table S1.**
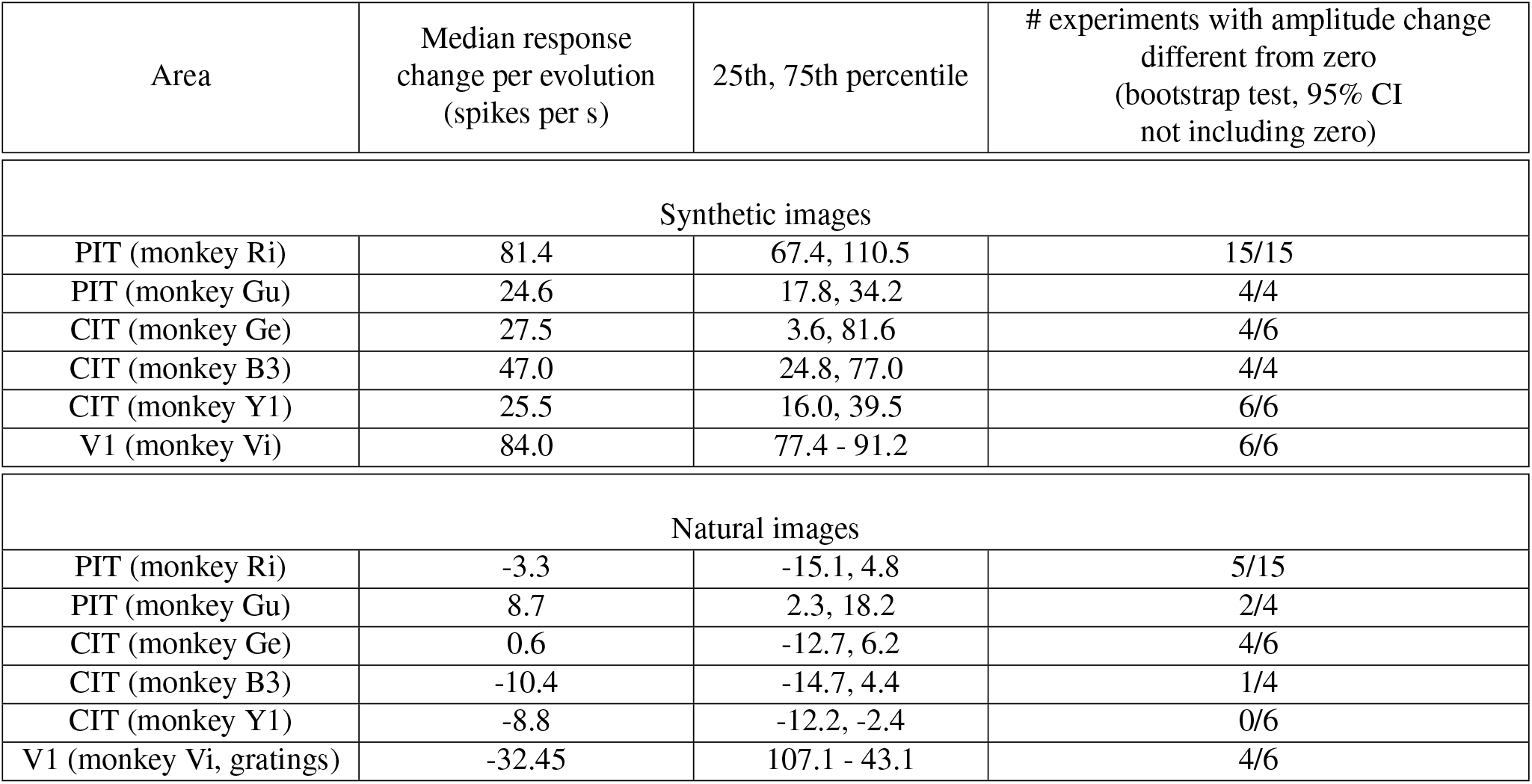
Response rate of neurons during evolution of synthetic images, across all experiments for each subject.

**Table S2.** Frequency that labeled ImageNet pictures matched evolved images for different animals (mean frequency ± standard error, per bootstrap).

|  | ImageNet labels |  |  |  |
| --- | --- | --- | --- | --- |
|  | ”macaque” | ”monkey” | ”face” (human only) | ”appliance” |
| frequency of label in sampled image set | $9.97 * 10^{-4}$ | $1.30 * 10^{-2}$ | $5.99 * 10^{-3}$ | $1.10 * 10^{-2}$ |
| Monkey Ri | 0.021±0.014 | 0.092±0.030 | 0.001±0.002 | 0.010±0.009 |
| Monkey Ge | 0.007±0.008 | 0.033±0.017 | 0.002±0.005 | 0.013±0.010 |
| Monkey B3 | 0.008±0.009 | 0.048±0.022 | 0.002±0.005 | 0.015±0.012 |
| Monkey Gu | 0.010±0.010 | 0.068±0.025 | 0.000±0.000 | 0.029±0.016 |
| Monkey Y1 | 0.002±0.005 | 0.041±0.017 | 0.001±0.003 | 0.041±0.019 |
|  |  | Probability that the values in Ri and Y1 were different: 0.070 |  | Probability that the values in Ri and Y1 were different: 0.076 |

**Table S3.**

| Subject (area) | Mean (spikes per s, ±sem) |  |  | Max (spikes per s, ±se) |  |  |
| --- | --- | --- | --- | --- | --- | --- |
|  | Synthetic | Reference (natural) | P < 0.03; Wilcoxon rank sum test, FDR correction | Synthetic | Reference (natural) | P < 0.03; Wilcoxon rank sum test, FDR correction |
| Ri (PIT) | 90.5±0.6 | 45.1±0.6 | 15 of 15 | 279.0±8.6 | 236.6±8.6 | 9 of 15<br>Synthetic > reference in 9/9 cases |
| Gu (PIT) | 26.6±0.4 | 21.3±0.4 | 3 of 4 | 122.4±4.1 | 121.4±4.6 | 0 of 4 |
| Ge (CIT) | 19.9±0.4 | -2.5±0.4 | 4 of 6 | 157.0±6.8 | 169.9±8.8 | 3 of 6<br>Synthetic > reference in 1/3 cases |
| B3 (CIT) | 45.0±0.4 | 5.9±0.3 | 4 of 4 | 213.1±4.9 | 169.9±18.2 | 3 of 4<br>Synthetic > reference in 3/3 cases |
| Y1 (CIT) | 34.0±0.4 | 14.5±0.4 | 6 of 6 | 156.4±8.9 | 146.3±6.7 | 1 of 6<br>synthetic > reference |
| Vi (V1) | 184.5±1.8 | 114.5±1.8 | 6 of 6 | 416.1±14.5 | (gratings)<br>390.3±13.0 | P values:<br>0.003, 0.003,<br>0.012, 0.050,<br>0.347 and 0.398 |

**Table S4.** Frequency that labeled ImageNet pictures matched evolved images for different animals (mean frequency ± standard error, per bootstrap).

| Mean firing rates to synthetic vs. top-prediction natural images (fc6-space experiments) |  |  |  | “Brute force” experiments, showing synthetic vs. >2,550 natural images in one session |  |  |  |
| --- | --- | --- | --- | --- | --- | --- | --- |
| Subject | Synthetic (mean across all experiments) | Natural images (close fc6 neighbors) | P value range across experiments | Subject | Natural (mean±sem, max±se, per bootstrap) | Synthetic | P value Wilcoxon rank sum test, permutation test(max) |
| Ri | 59.4±1.4<br>N = 4 | 30.8±1.3 | $4.5 * 10^{-144}$<br>to $3.5 * 10^{-8}$ | Ri | 24.7±0.5<br>104.2±1.4 | 72.3±1.9<br>130.3±5.8 | $< 1 * 10^{-6}$<br>$1.0 * 10^{-3}$ |
| Gu | 38.5±0.8<br>N = 3 | 26.3±1.4 | $5.8 * 10^{-309}$<br>to $5.1 * 10^{-2}$ | Ge | -8.4<br>87.0±3.8 | 28.0<br>83.5±4.4 | $< 1 * 10^{-6}$<br>$1.0 * 10^{-3}$ |
| Y1 | 38.3±1.1<br>N = 3 | 21.4±2.2 | $8.7 * 10^{-23}$<br>to $1.0 * 10^{-2}$ | - | - | - | - |

